# Genetic Characteristics and Phylogeny of 969-bp S Gene Sequence of SARS-CoV-2 from Hawaii Reveals the Worldwide Emerging P681H Mutation

**DOI:** 10.1101/2021.01.06.425497

**Authors:** David P. Maison, Lauren L. Ching, Cecilia M. Shikuma, Vivek R. Nerurkar

**Author notes:** **Corresponding Author:** Vivek R. Nerurkar, Ph.D., Department of Tropical Medicine, Medical Microbiology and Pharmacology, John A. Burns School of Medicine, 651 Ilalo Street, BSB 320, Honolulu, Hawaii 96813.

## Abstract

COVID-19 pandemic has ravaged the world, caused over 1.8 million deaths in the first year, and severely affected the global economy. Hawaii is not spared from the transmission of SARS-CoV-2 in the local population, including high infection rates in racial and ethnic minorities. Early in the pandemic, we described in this journal various technologies used for the detection of SARS-CoV-2. Herein we characterize a 969-bp SARS-CoV-2 segment of the S gene downstream of the receptor-binding domain. At the John A. Burns School of Medicine Biocontainment Facility, RNA was extracted from an oropharyngeal swab and a nasal swab from two patients from Hawaii who were infected with the SARS-CoV-2 in August 2020. Following PCR, the two viral strains were sequenced using Sanger sequencing, and phylogenetic trees were generated using MEGAX. Phylogenetic tree results indicate that the virus has been introduced to Hawaii from multiple sources. Further, we decoded 13 single nucleotide polymorphisms across 13 unique SARS-CoV-2 genomes within this region of the S gene, with one non-synonymous mutation (P681H) found in the two Hawaii strains. The P681H mutation has unique and emerging characteristics with a significant exponential increase in worldwide frequency when compared to the plateauing of the now universal D614G mutation. The P681H mutation is also characteristic of the new SARS-CoV-2 variants from the United Kingdom and Nigeria. Additionally, several mutations resulting in cysteine residues were detected, potentially resulting in disruption of the disulfide bridges in and around the receptor-binding domain. Targeted sequence characterization is warranted to determine the origin of multiple introductions of SARS-CoV-2 circulating in Hawaii.

## Introduction

The zoonotic virus responsible for the present Coronavirus Disease 2019 (COVID-19) pandemic is SARS-CoV-2 (formerly nCoV).^1^ SARS-CoV-2 emerged in Wuhan, China,^2^ at a seafood market^3^ in November 2019^4^ and has been evolving ever since.^1^ The COVID-19 pandemic resultant from SARS-CoV-2 has been responsible for infecting more than 84 million people worldwide and has been fatal in more than 1.8 million persons experiencing an infection.^5^ The State of Hawaii has dealt with more than 22,000 cases and 280 deaths, with daily reports steady at approximately 60 new cases per day since August 2020.^6^

To understand SARS-CoV-2 emergence and the disease, COVID-19, one must look at the genome, as the virus evolves and emerges through genomic alterations and adaptations. SARS-CoV-2 belongs to the family of *Coronaviridae* – genus Betacoronavirus - which are viruses with 26,000-32,000 nucleotide long single-stranded positive-sense RNA genomes.^7–9^ Geneticists and virologists look at the SARS-CoV-2 genome and its adaptations to analyze the nucleotide and amino acid variations. Analysis of the SARS-CoV-2 genome will allow us to track the spread through unique genomic fingerprints,^1^ determine whether these adaptations alter the viral fitness, infectious capabilities,^10^ and develop potential vaccines and therapeutics.^11,12^

While genes encode 20 proteins consisting of four structural and 16 non-structural proteins in the SARS-CoV-2 ~30,000-bp genome,^9^ the gene looked to most is the S gene responsible for the spike protein. The spike protein is a 1,273 amino acid long^12^ (YP_009724390.1)(NC_045512) surface protein that is the viral component accountable for interacting with the human ACE2 (UNIPROT ID Q9BYF1).^7,13,14^ ACE2, genetically encoded on the human X chromosome (ENSG00000130234), is a component of the renin-angiotensin hormone system in humans and is ultimately a vasodilator.^15,16^ This interaction between SARS-CoV-2 spike protein and human ACE2 via the RBD allows SARS-CoV-2 to enter cells and infect the human host.^12^ Mutations in the spike protein can alter binding efficiency and viral fitness.^10^ Indeed, some nucleotide mutations in the SARS-CoV-2 S gene change pathogenicity^12^ and fitness,^10,17^ reduce virulence,^1,4^ and have become commonplace in tracking the spread of SARS-CoV-2.^17^ These S gene and spike protein mutations can increase transmission of the virus between hosts through anatomical localization to the upper respiratory tract.^10^ Therefore, to understand the SARS-CoV-2 pathogenicity, it is important to characterize the virus mutations to study viral pathogenesis and vaccine development. In this study we report analysis of a 969-bp SARS-CoV-2 S gene from two patients from Hawaii to understand the changes in the spike protein, a target for vaccines.

## Methods

### Patient Samples and Viral RNA Extraction

Two patients (PID 00498 and PID 00708) analyzed in this report were part of the University of Hawaii at Manoa IRB approved H051 study (IRB# 2020-00367) led by Dr. Cecilia Shikuma. Patient PID 00498 and patient PID 00708 are both males, one is caucasian and the other is Japanese/Okinawan/Filipino, and their mean age is 29.5 years. Oropharyngeal swab (OS - PID 00498) and Nasal swab (NS - PID 00708) from previously identified SARS-CoV-2 positive patients were collected and stored at −80°C. Swabs from PID 00498 and from PID 00708, were collected in August 2020, 3 days after first PCR positive diagnosis. Neither of the patients traveled outside of Honolulu in the week before their first PCR positive SARS-CoV-2 diagnosis, however both identified potential sources of exposure in Honolulu.

Swabs stored in VTM at −80C were thawed in the biosafety cabinet at the John A. Burns School of Medicine high containment laboratory as part of the University of Hawaii at Manoa IBC approved study (IBC#20-04-830-05). VTM was centrifuged to separate the supernatant from debris, aliquoted and 140 μL of the VTM was used for viral RNA purification using the QIAamp^®^ Viral RNA Mini Kit (Cat# 52906) following the manufacturer’s instructions. The samples were eluted in 30 μL of the elution buffer.

### Reverse-Transcriptase Polymerase Chain Reaction and Sequencing

As per the manufacturer’s instructions, purified viral RNA was transcribed into cDNA using the Takara *LA Taq* polymerase Kit (Cat #RR012A) with random nine-mers and extension time of 90 minutes. Primer sets were designed based on published sequences and were procured from Integrated DNA Technologies (Coralville, IA). A 1,127-bp segment of the S gene was amplified using the Takara RNA LA PCR Kit (Cat #RR012A) and primers CF and CR (Figure 1).^18^ PCR was conducted according to the manufacturer’s instructions and cycled on the Applied Biosystems GeneAmp^®^ PCR System 9600. PCR products were then electrophoresed on 1.5% agarose 1x TBE gels at 50V and the amplicons of interest were purified using the Qiagen QIAquick Gel Extraction Kit (Cat# 28704).

**Figure 1:**
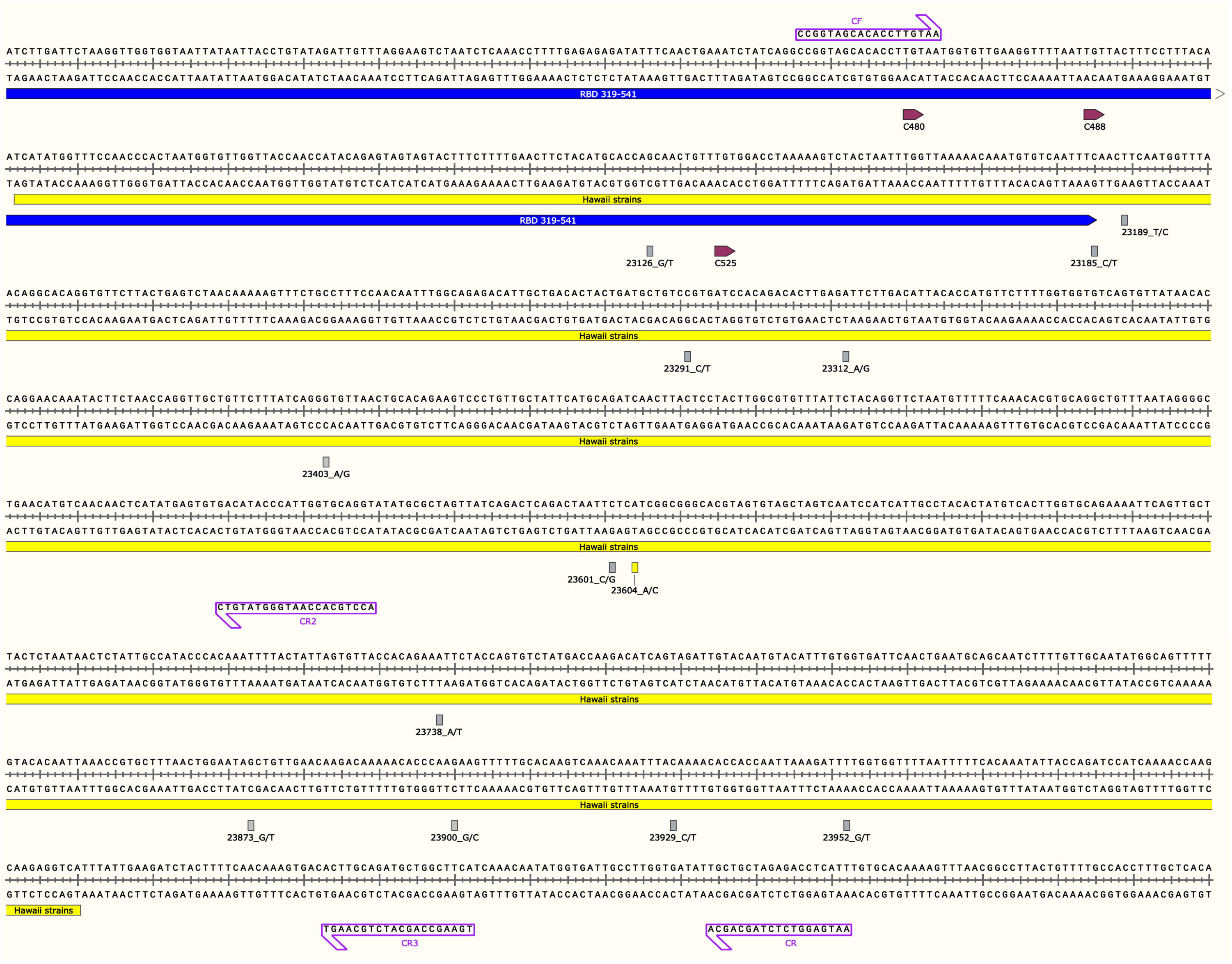
SARS-CoV-2 S Gene Region used in this Study Along with Annotated Primers, Mutations, and Cysteine Residues of the Receptor Binding Domain. Figure represents the Hawaii strain MW237663 and MW237664 sequences. The primer pair, CF/CR, was used to amplify the 1,127-bp S gene fragment and primers CF, CR, CR2 and CR3 depicted with purple boxes were used for Sanger sequencing. The yellow box indicates the start and end of the 969-bp sequence. The blue line indicates the 3’ end of the S gene receptor binding domain (RBD). RBD cysteine residues are shown in depicted boxes. All mutations found in this study are in their respective loci with nucleotide numbers and rectangular boxes correlating to the sense strand as indicated after the nucleotide number and underscore in the figure (nucleotide/protein mutations: G23126T/A522S, C23185T/F541F, T23189C/F543L, C23291T/R577C, A23312G/I584V, A23403G/D614G, C23601G/S680C, C23604A/P681H, A23738T/I726F, G23873T/A771S, G23900C/E780Q, C23929T/Y790Y, and T23952G/F797C). All boxes are grey except for the P681H mutation seen in the Hawaii strains from this study, shown with a yellow rectangle. Image was generated with the SnapGene software (from Insightful Science; available at snapgene.com) and was created with BioRender.com.

Sanger sequencing was conducted on the amplicons using four primers (CF, CR,^18^ CR2, and CR3) at the ASGPB core facility at the University of Hawai□i at Mānoa (Figure 1). The resulting sequences were input into and verified using both MEGAX^19,20^ and SnapGene software (Insightful Science, snapgene.com) and aligned using MUSCLE program^21^ to define the contiguous sequence. The resulting 969-bp consensus sequences were uploaded to NCBI. The S gene 969-bp region encompasses nucleotides 23,042 to 24,010, and corresponds to amino acids 494 to 816, which involves the 3’ and C-terminal of the RBD that ends at nucleotide 23,185 and amino acid 541.^7^

### SNP Analysis

Sixty-eight coronavirus strains representing alpha and beta lineages were manually selected from NCBI. Of these 68 strains, 55 were SARS-CoV-2 strains which represented 25 distinct geographical locations spanning the pandemic duration, at least one per month from December 2019 to September 2020. All SARS-CoV-2 sequences, including previously published Hawaii sequences, were first aligned and redundant sequences were removed from further analysis. Coronavirus sequences were aligned with the 969-bp S gene region with SnapGene using MUSCLE and the corresponding region was used for future analysis.^21^ The non-SARS-CoV-2 strains were removed if the 969-bp S gene of SARS-CoV-2 sequence did not align with the S gene of the non-SARS-CoV-2 strains. Based on the alignment SNPs were identified and annotated into SnapGene to analyze the amino acid substitutions.

Upon finding the P681H mutation among the two Hawaii strains in this study, the GISAID database^22,23^ was used to filter worldwide SARS-CoV-2 sequences by the P681H mutation to create a ratio of sequences containing the P681H mutation to all sequences reported in the GISAID database for a given month. Inclusion criteria were for sequences providing a full month, day, and year. The D614G mutation underwent assessment in the same manner for comparison. All prevalence data converted into ratio underwent a logarithmic transformation. Pearson’s correlation tests between P681H frequency vs. month, D614G frequency vs. month, and P681H frequency vs. D614G frequency were conducted and verified using GraphPad Prism version 9.0.0 for Mac (GraphPad Software, San Diego, California USA, www.graphpad.com), JASP version 0.14,^24^ and RStudio version 1.3.1093.^25^

### Phylogenetic Tree

After the SNP analysis, incomplete sequences were removed prior to the construction of the phylogenetic tree. The phylogenetic tree was constructed using MEGAX.^19^ The alignment was first done using the program MUSCLE.^21^ The phylogenetic tree was then generated with Maximum Likelihood parameters with 1,000 bootstraps in MEGAX^19,20^ using the University of Hawaii MANA High Performance Cluster. The output tree from MEGAX was rooted using FigTree version 1.4.4 based on alpha coronavirus human 299E (KF514433).^26^

## Results

### Gene Amplification and Sequence Analysis

SARS-CoV-2 genomic sequences were detected by RT-PCR in both the patients, PID 00498 and PID 00708, using various primers spanning the S gene (Figure 1). A 1,127-bp segment was amplified and sequenced and the entire sequence of the amplicon was aligned with at least one forward and one reverse sequence (translated into reverse-complement) to span the whole 1,127-bp region. For final sequence analysis a 969-bp sequence verified by sequencing the 5’ and 3’ ends was used. The two Hawaii sequences were deposited in the GenBank, accession numbers MW237663 for PID 00498 and MW237664 for PID 00708.

### SNP Analysis

Of the 55 original non-Hawaii SARS-CoV-2 strains, 47 were redundant in the 969-bp segment of the S gene. Of the 12 Hawaii strains deposited in the GenBank, nine were redundant in the 969-bp sequence region. Thus, we analyzed eight non-Hawaii SARS-CoV-2 strains and three SARS-CoV-2 strains from Hawaii. With the addition of the two SARS-CoV-2 strain sequences from this study, a total of 13 SARS-CoV-2 sequences were compared and SNPs encompassing the 969-bp region of the S gene were analyzed (Table 1). The alignment containing the 13 sequences revealed 13 SNPs (Table 1)(Figure 1). Eleven of the thirteen mutations resulted in non-synonymous mutations (A522S, F543L, R577C, I584V, D614G, S680C, P681H, I726F, A771S, E780Q, and F797C) (Table 1 and Figure 1). Two of the thirteen mutations resulted in a synonymous mutation (amino acid 541 and 790) (Table 1 and Figure 1). The P681H mutation is unique to the Hawaii strains from this study (MW237663 and MW237664).

**Table 1:**
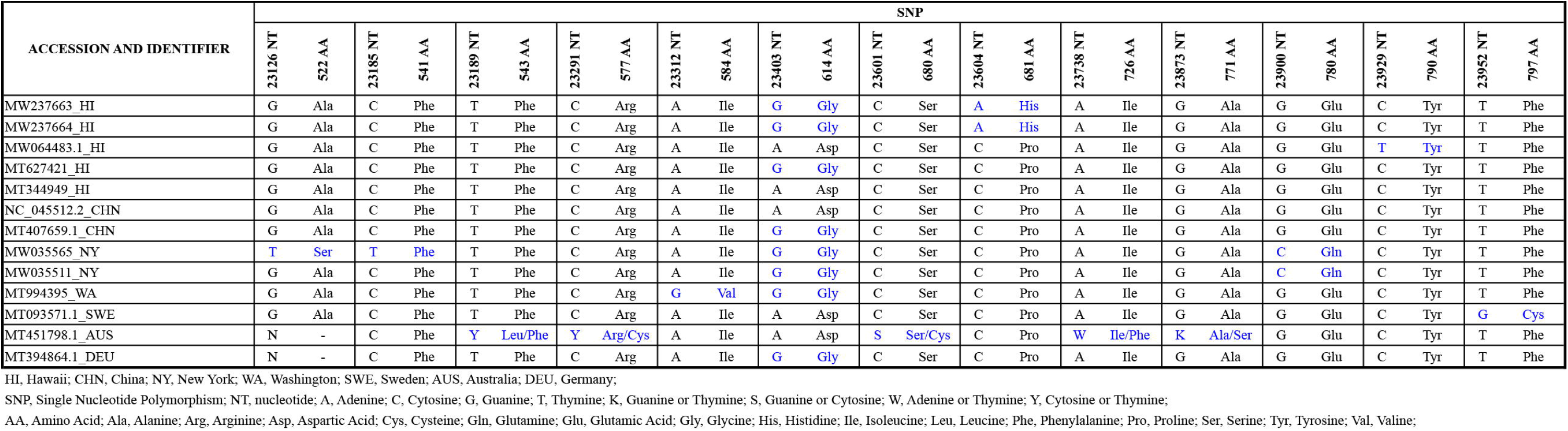
Comparison of Single Nucleotide Polymorphisms and Resultant Amino Acid Substitutions in a 969-bp S gene of SARS-CoV-2 Strains from Hawaii

GISAID reported the first P681H mutation on March 12, 2020 (EPI_ISL_430887).^27^ Further, from March 01, 2020 through December 31, 2020 GISAID reports a total of 5,955 strains that have the P681H mutation. During that same time, GISAID has reported approximately 296,064 SARS-CoV-2 strains (Table 2). Pearson’s correlation between time in months versus prevalence of P681H (Figure 2A) and D614G (Figure 2B) of logarithmically transformed data indicates an increase in the number of strains having the P681H mutation (*r* = 0.96, *P* < 0.0001) (Figure 2A) and plateauing of the D614G mutation (*r* = 0.78, *P* = 0.008) (Figure 2B). P681H mutations were not reported in May 2020. Further, Pearson’s correlation indicates positive correlation (*r* = 0.71, *P* = 0.03) between the worldwide prevalence of P681H and D614G (Figure 2C).

**Figure 2:**
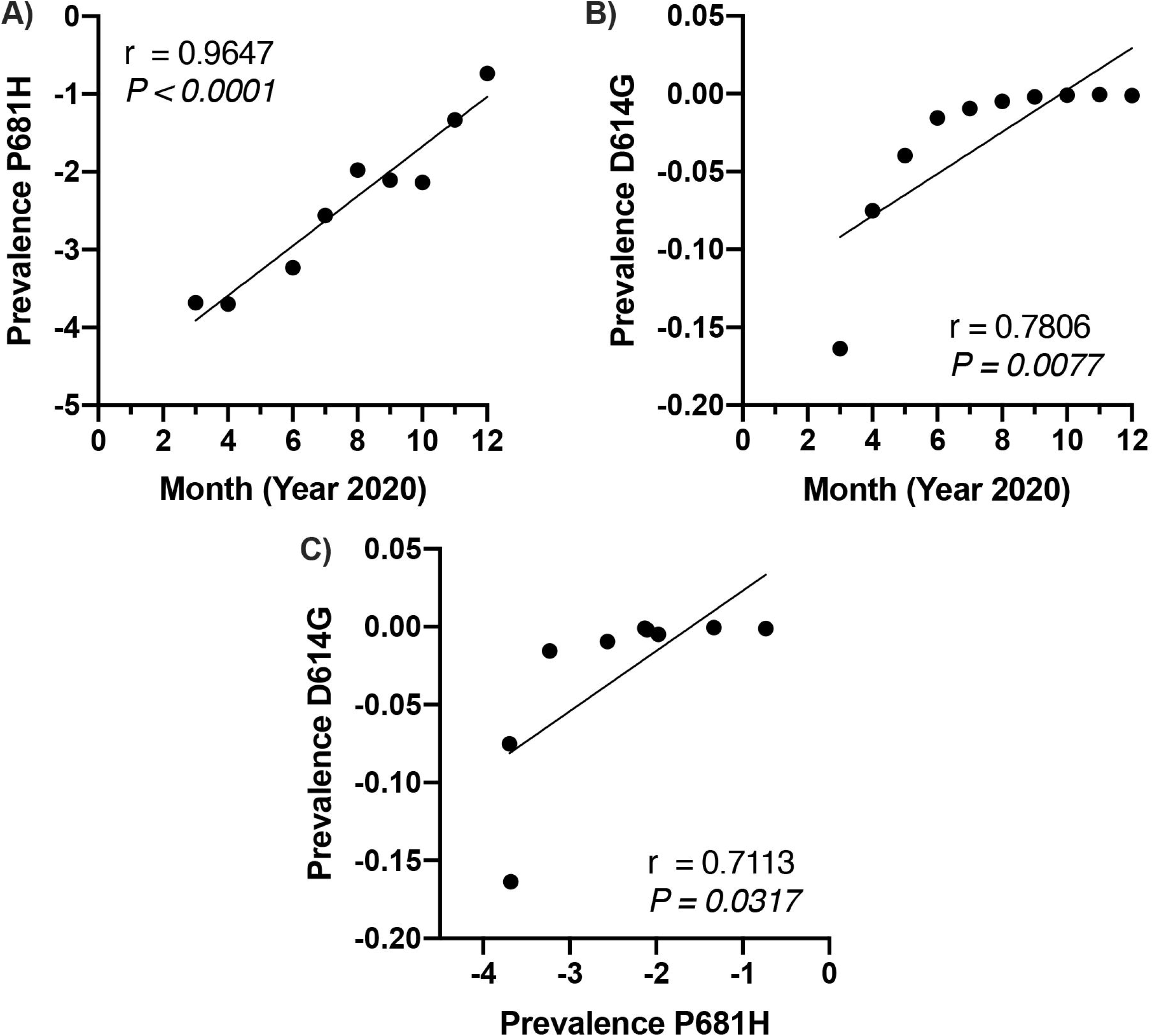
Pearson’s Correlation of Logarithmically Transformed Data Showing Positive Correlation for Time in Months Versus Both P681H and D614G Mutations. A) Graphical representation of the logarithmic transformed ratio of P681H mutation among all reported GISAID strains on the y-axis and month on the x-axis (ex: 3 = March, 4 = April, etc.). Linear regression line shown along with Pearson’s correlation, r = 0.97, *P* = 0.00003. B) Graphical representation of the logarithmic transformed ratio of D614G mutation among all reported GISAID strains on the y-axis and month on the x-axis. Linear regression line shown along with Pearson’s correlation, r = 0.78, *P* = 0.008. C) Graphical representation of the logarithmic transformed ratio of D614G mutation among all reported GISAID strains on the y-axis and the logarithmic transformed ratio of P681H mutations among all reported GISAID strains on the x-axis. Linear regression line shown along with Pearson’s correlation, r = 0.71, *P* = 0.03. Graphs created with GraphPad Prism version 9.0.0 for Mac (GraphPad Software, San Diego, California USA, www.graphpad.com).

**Table 2:**
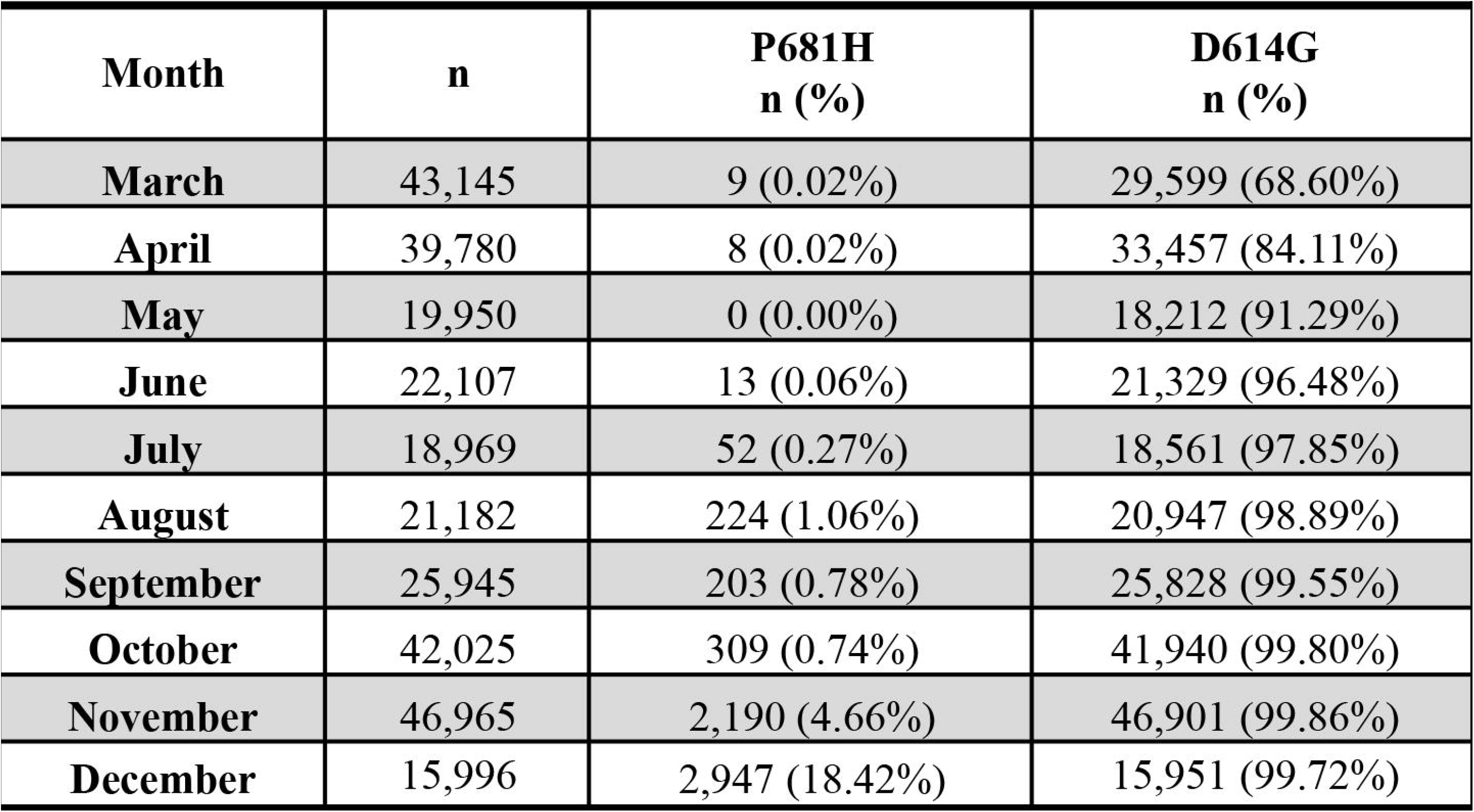
The Distribution and Frequency of P681H and D614G Mutations Among All SARS-CoV-2 Sequences by Month Reported in the GISAID Database in 2020

| Table 2. The Distribution and Frequency of P681H and D614G Mutations Among All SARS-CoV-2 Sequences by Month Reported in the GISAID Database in Year 2020 |  |  |  |
| --- | --- | --- | --- |
| Month | n | P681H<br>n (%) | D614G<br>n (%) |
| March | 43,145 | 9 (0.02%) | 29,599 (68.60%) |
| April | 39,780 | 8 (0.02%) | 33,457 (84.11%) |
| May | 19,950 | 0 (0.00%) | 18,212 (91.29%) |
| June | 22,107 | 13 (0.06%) | 21,329 (96.48%) |
| July | 18,969 | 52 (0.27%) | 18,561 (97.85%) |
| August | 21,182 | 224 (1.06%) | 20,947 (98.89%) |
| September | 25,945 | 203 (0.78%) | 25,828 (99.55%) |
| October | 42,025 | 309 (0.74%) | 41,940 (99.80%) |
| November | 46,965 | 2,190 (4.66%) | 46,901 (99.86%) |
| December | 15,996 | 2,947 (18.42%) | 15,951 (99.72%) |

### Phylogenetic Tree

Of the 13 SARS-CoV-2 sequences used for SNP identification, two were incomplete due to unidentified nucleotides and were removed from the phylogenetic analysis. Similarly, of the 13 non-SARS-CoV-2 sequences, four did not align to the 969-bp S gene due to large insertions and/or deletions and were removed from further analysis. Therefore, the final phylogenetic tree was constructed using 20 coronavirus sequences, 11 SARS-CoV-2 and nine non-SARS-CoV-2 sequences (Figure 3). Based on the phylogenetic tree constructed using the Maximum Likelihood method, the alpha and beta coronaviruses segregated as expected. Further, the beta coronaviruses lineages A, B, C and D segregated with a bootstrap value of >70. Within the beta coronavirus lineage B, the SARS-CoV-1 and the bat coronaviruses were distinctly segregated from the SARS-CoV-2 with a bootstrap value of 92.

**Figure 3:**
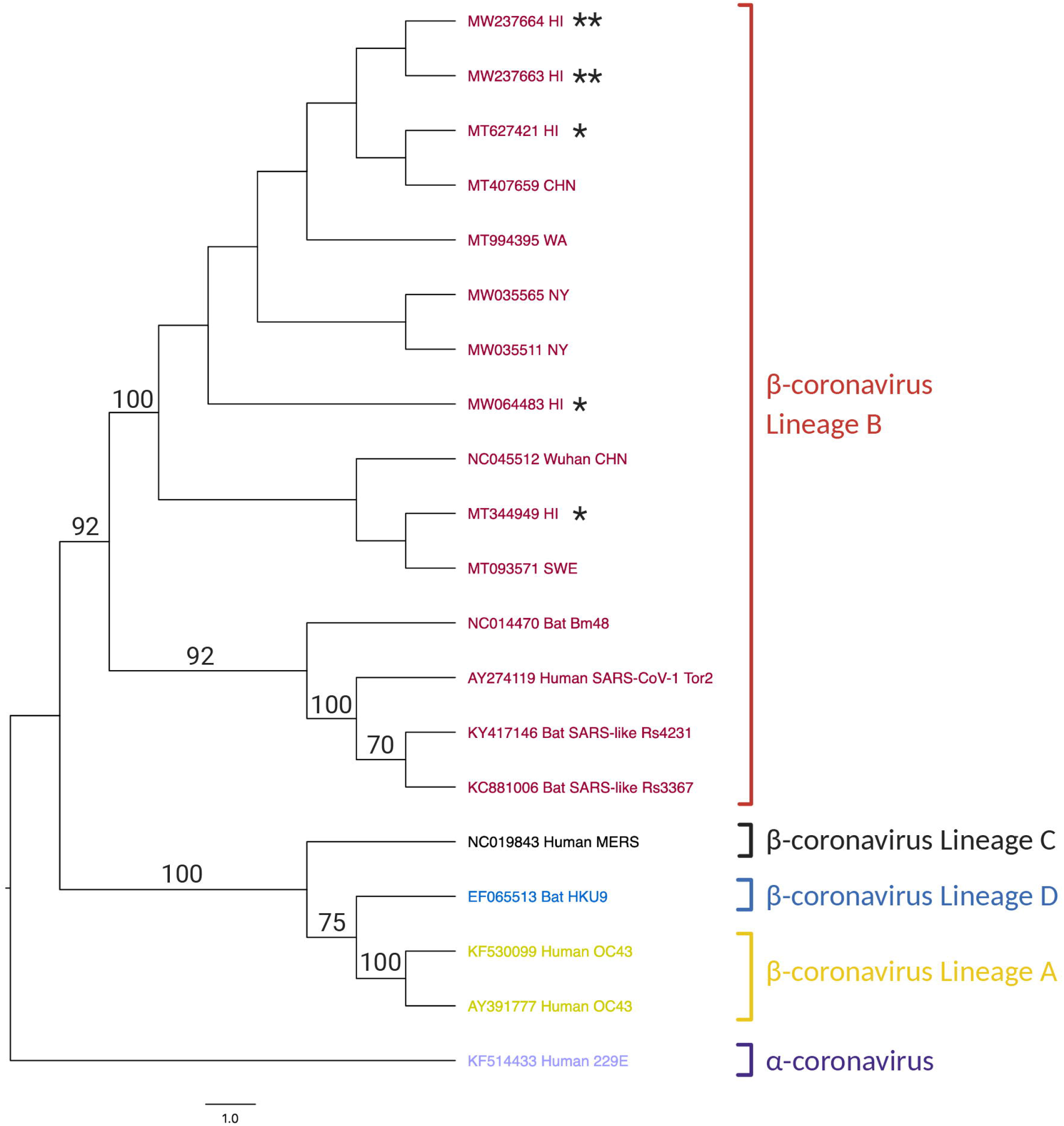
Maximum Likelihood Phylogenetic Tree Constructed Using a 969-bp S gene Region of SARS-CoV-2. The Tamura-Nei model and the Maximum Likelihood method were used to infer evolutionary history using 1,000 bootstraps. The tree with the highest log likelihood is shown in the figure. Next to the branches is shown the percentage of trees in which the associated taxa clustered together - only values greater than 70 are displayed. Neighbor-Join and BioNJ algorithms were applied to a matrix of pairwise distances estimated using the Tamura-Nei model to automatically obtain the initial tree for the heuristic search, and then selecting the topology with superior log likelihood value. This analysis involves nucleotide sequences from 20 taxa. There were a total of 1,366 sites. Evolutionary analyses were conducted in MEGAX^19,20^ using the University of Hawaii MANA High-Performance Cluster. The tree was rooted using the alphacoronavirus human 229E (KF514433) taxa in FigTree version 1.4.4. Yellow text is used for betacoronavirus lineage A, red text denotes betacoronavirus lineage B, black text denotes betacoronavirus lineage C, blue text shows betacoronavirus lineage D, and purple text is for alphacoronavirus. * represents strains from Hawaii. ** represents strains from this study. Created with BioRender.com.

Within the SARS-CoV-2 branch of beta coronavirus lineage B, the D614G mutation was the defining node for branch separation. The three sequences lacking the D614G mutation (NC_045512, MT344949, and MT093571) are separated from sequences with the D614G mutation with a bootstrap value of 100, except for the MW066483 Hawaii sequence, which also does not have the D614G mutation. Wuhan (NC_045512) and Hawaii (MT344949) strains are identical and contain no mutations, as NC_045512 is the reference genome for SARS-CoV-2. The Sweden strain (MT093571) has a F797C mutation. The Hawaii strain MW064483 is the next closest cluster to the D614G defining node and additionally contains the synonymous tyrosine mutation at amino acid 790. All the remaining sequences have the D614G mutation. Clustering near the Hawaii Strain MW066483 are the two strains from New York, USA (MW035565 and MW035511), both containing the E780Q mutation and MW035565 also containing the A522S mutation and a synonymous phenylalanine mutation at amino acid 541. Branching from the New York strains cluster is the Washington, USA, strain (MT994395), exclusively having the I584V mutation. The two Hawaii strains from this study (MW237663 and MW237664) have the emerging P681H mutation and cluster closely with previously published SARS-CoV-2 sequences from Hawaii (MT627421) and China (MT407659). MT627421 strain from Hawaii and MT407659 strain from China are identical and are the only sequences to contain the D614G mutation exclusively.

In summary, SARS-CoV-2 strains from Hawaii deposited in the GenBank in March 2020 clustered with sequences from Wuhan, Sweden, China, and the state of New York, USA. The SARS-CoV-2 strains in this study cluster from the state of Washington, USA and with sequences from China and Hawaii. Five of the 13 SARS-CoV-2 sequences used in the phylogenetic tree are sequences from Hawaii, marked with asterisk(s). Coronavirus lineage determinations are based on phylogenetic trees constructed by Chan and colleagues, 2020,^28^ and Su and colleagues, 2016.^29^

## Discussion

This report focuses on SARS-CoV-2 strains from two patients in Hawaii based on a 969-bp S gene. The sequence and phylogenetic analysis indicate and support that the S gene is continuously mutating as previously reported,^30^ and Hawaii may be harboring a unique strain with an emerging mutation in an altered spike protein. Analysis of the SNPs found in the 969-bp S gene region, indicates the loss of a proline residue and the gain of cysteine residues. These mutations potentially alter the spike protein monomeric and trimeric structures.

The P681H represents the loss of a proline residue and the gain instead of an imidazole-containing histidine residue. According to the GISAID database, the P681H mutation is found worldwide in 5,955 strains reported as of December 31, 2020. The Pearson’s correlation test of the logarithmic transformed P681H prevalence of the mutation versus time indicates that the P681H mutation is exponentially increasing worldwide and the sequences encompassing the P681H mutation is dominating significantly when compared to other SARS-CoV-2 strains. Such a significant finding indicates that there is a selective pressure in favor of this mutation.

A study looking at the effect of the loss of a proline residue in the spike protein of the mouse hepatitis virus, a coronavirus, observed altered pathology, fusion kinetics, and enhanced infectivity.^31^ The same study also suggests that prolines in regions adjacent to RBDs may not be essential for fusion but may significantly change structure and function of the spike protein.^31^ Recent SARS-CoV-2 studies indicate that the P681H mutation is immediately juxtaposed to the amino acid 682-685, furin cleavage site, identified at the S1/S2 linkage site, which predicted enhance systemic infection^12,32^, and increased membrane fusion.^11^ Additionally, the proline in the P681H mutation is within the epitope found to be the highest-ranking B and T cell epitope based on the in silico long-term population-scale epitope prediction for vaccine development study.^11^ That is, the sequence surrounding this P681H mutation is predictably the loci where the immune response is targeted. Meaning this mutation could be the first identified SARS-CoV-2 mutation of antigenic evolution.^1^ Further studies are warranted to analyze the pathogenicity and virulence of the newly identified P681H mutation seen in the Hawaii strains and whether this is a viral evasion mechanism to deter antibody recognition or another increase in fitness. Now that SARS-CoV-2 vaccines are available in the United States under the FDA-EUA, it is critical to evaluate the epitope-altering mutations. These mutations could change the effectiveness of FDA-EUA SARS-CoV-2 vaccines that rely on the structure of the spike protein.^33,34^

Twelve full length sequences have been published from Hawaii (MT627420.1, MT627421.1, MW064481.1, MW064482.1, MW064483.1, MW064596.1, MW064825.1, MW064826.1, MW065225.1, MW190887.1, MT344948.1, MT344949.1). All of these original strains introduced to Hawaii were collected in March 2020, and 66.6% (8/12) have the D614G mutation. The D614G mutation has become universal throughout the SARS-CoV-2 strains.^10^ The D614G mutation is known to enhance infectivity, replication, and localize the virus to the upper respiratory tract to increase transmission.^10,17^ Interestingly, several mutations (nucleotide position 241 C→T, position 3037 C→T, and position 14408 C→T, etc.) exist alongside the D614G mutation^10,17^ Similarly, in the both Hawaii SARS-CoV-2 strains the P681H mutation is also present alongside D614G^10,13,17^. This observation indicates that, similar to D614G mutation the P681H mutation is becoming globally prevalent among SARS-CoV-2 sequences.

The R577C, S680C, and F797C mutations depicted in Table 1 are also very prominent mutations in that they present possible new disulfide bridges forming within and around the RBD. The RBD possesses eight cysteine residues, with disulfide bridges formed between amino acids 336:361, 379:432, 480:488, and 391:525.^7,14^ The aforementioned cysteine mutations may interact with these known bridges or create new bridges. The F797C mutation is seen alone in the Sweden strain (MT093571.1). The R577C and S680C mutations are present together in the strain from Australia (MT451798.1). Similar to the P681H, the S680C mutation is also within the epitope region of the B and T cell epitope in silico prediction model for vaccine development.^11^ Further studies are warranted to evaluate the disulfide bridge configurations and whether an odd number of cysteines in this region can result in a dynamic bridge or the addition of several cysteines can alter the spike protein structure. Such studies would help to understand if these mutations are evolutionary mechanisms that alter virulence, or perhaps influence fusion kinetics. The other non-synonymous SNPs found in this study - A522S, F543L, I584V, I726F, A771S, E780Q - are not as apparent in presenting drastic evolutionary change, but they too deserve further analysis.

Recently, the NERVTAG group based out of London, England, has reported on a new SARS-CoV-2 variant (VOC202012/01).^35,36^ NERVTAG reports that the variant has increased transmissibility and further studies are underway to confirm their report.^37^ This variant includes amino acid mutations in ORF1ab, spike, Orf8, and N.^35,37,38^ The six ORF1ab mutations are T1001I, A1708D, I2230T, and ΔS3675, ΔG2676, and ΔF3677.^35,37,38^ The ten spike mutations are ΔH69, ΔV70, ΔY145, N501Y, A570D, D614G, P681H, T716I, S982A, and D1118.^35,37,38^ The Orf8 mutations are Q27stop, R52I, and Y73C.^37,38^ The N protein mutations are D3L and S235F.^37,38^ When comparing the SNPs encompassing the 969-bp of two strains from this study to the reference genome for VOC202012/01, EPI_ISL_601443,^39^ we found two similar mutations, D614G and P681H.^37^ Further, EPI_ISL_601443 shows the N501Y, A570D, D614G, P681H, and T716 mutations in the 969-bp region, while the two Hawaii strains, MW237663 and MW237664, display the D614G and P681H mutations. Additionally, a new variant in Nigeria (B.1.207)(EPI_ISL_729975)^40^ has been defined by the P681H mutation found in the two Hawaii strains.

The two Hawaii strains analyzed in this study cluster together predictably due to the emerging P681H mutation. These two strains also cluster closely with a strain from China and a previously published Hawaii strain. Other previously published Hawaii strains cluster with SARS-CoV-2 strains from New York, Wuhan, China, and Sweden. These analysis and resultant phylogenetic tree indicate that the virus has likely been introduced to Hawaii through several sources.

Over the past year, SARS-CoV-2 worldwide has evolved and will continue to do so. As of this report’s submission, four new SARS-CoV-2 variants have been reported from the United Kingdom (VOC 202012/01/B.1.1.7),^41^ South Africa (501Y.V2),^41^ Denmark (Mink Cluster V),^42,43^ and Nigeria (B.1.207).^41^ This fast pace of evolutionary changes will affect pathogenicity of SARS-CoV-2^12,44^ and warrants further in silico, in vitro, and in vivo studies.

In summary, COVID-19 in Hawaii and the pandemic originating in Wuhan in the 2019-2020 winter is still ongoing. The virus continues to mutate and the effects and outcomes of several of these mutations has yet to be elucidated. This study demonstrates a partial sequence from the first SARS-CoV-2 strain possessing the P681H non-synonymous mutation. In Hawaii, Native Hawaiians and Pacific Islanders have significantly high prevalence of SARS-CoV-2 when compared to other ethnic minorities and Whites. Characterizing viral sequences from these minority groups is important to better understand virus transmission and pathogenicity.

## Acknowledgements

This research was supported by a grant (P30GM114737) from the Pacific Center for Emerging Infectious Diseases Research, COBRE, National Institute of General Medical Sciences, NIH; by a grant (U54MD007601) from Ola Hawaii, National Institute on Minority Health and Health Disparities, NIH; and by a contract (CT-MAY-2000282) from the City and County of Honolulu. We thank Dr. Sean Cleveland, Information Technology Services, University of Hawaii (UH) for assistance with the phylogenetic tree and the University of Hawaii MANA High-Performance Cluster for use of the facility, and Dr. Vedbar Khadka for assistance with MEGAX. We thank the Nurses and staff of the Hawaii Center for AIDS for assisting with the H051 study, and the patients for participating in this study.

## Author’s Contributions

- Guarantor of integrity of entire study: David P. Maison, Vivek R. Nerurkar
- Study concept design: Vivek R. Nerurkar, David P. Maison
- Data acquisition/analysis: David P. Maison, Lauren L. Ching, Vivek R. Nerurkar
- Manuscript drafting/revision for intellectual content: David P. Maison, Vivek R. Nerurkar
- Literature review: David P. Maison, Vivek R. Nerurkar
- Clinical studies: Cecilia M. Shikuma
- Statistics: David P. Maison, Vivek R., Nerurkar
- Manuscript editing: David P. Maison, Lauren L. Ching, Cecilia M. Shikuma, Vivek R. Nerurkar

## Abbreviations

ASGPB: Advanced Studies in Genomics, Proteomics, and Bioinformatics
A1708D: Alanine to Aspartic Acid at Amino Acid 1708
A570D: Alanine to Aspartic Acid at Amino Acid 570
A522S: Alanine to Serine at Amino Acid 522
A771S: Alanine to Serine at Amino Acid 771
ACE2: Angiotensin-Converting Enzyme 2
R577C: Arginine to Cysteine at Amino Acid 577
R52I: Arginine to Isoleucine at Amino Acid 52
N501Y: Asparagine to Tyrosine at Amino Acid 501
D614G: Aspartic Acid to Glycine at Amino Acid 614
D1118H: Aspartic Acid to Histidine at Amino Acid 1118
D3L: Aspartic Acid to Leucine at Amino Acid 3
cDNA: complementary deoxyribonucleic acid
COVID-19: Coronavirus Disease 2019
ΔG2676: deletion of Glycine Amino Acid 2676
ΔH69: deletion of Histidine Amino Acid 69
ΔF3677: deletion of Phenylalanine Amino Acid 3677
ΔS3675: deletion of Serine Amino Acid 3675
ΔY145: deletion of Tyrosine Amino Acid 145
ΔV70: deletion of Valine Amino Acid 70
DNA: deoxyribonucleic acid
EUA: Emergency Use Authorization
GISAID: Global Initiative of Sharing All Influenza Data
E780Q: Glutamic Acid to Glutamine at Amino Acid 780
Q27stop: Glutamine to stop codon at Amino Acid 27
IBC: Institutional Biosafety Committee
IRB: Institutional Review Board
I726F: Isoleucine to Phenylalanine at Amino Acid 726
I2230T: Isoleucine to Threonine at Amino Acid 2230
I584V: Isoleucine to Valine at Amino Acid 584
MUSCLE: Multiple Sequence Comparison by Log-Expectation
NCBI: National Center for Biotechnology Information
NERVTAG: New and Emerging Respiratory Virus Threats Advisory Group
nCoV: novel coronavirus
PID: Patient Identification
F797C: Phenylalanine to Cysteine at Amino Acid 797
F543L: Phenylalanine to Leucine at Amino Acid 543
PCR: Polymerase Chain Reaction
P681H: Proline to Histidine at Amino Acid 681
RBD: Receptor Binding Domain
RT-PCR: reverse transcriptase Polymerase Chain Reaction
RNA: ribonucleic acid
S982A: Serine to Alanine at Amino Acid 982
S680C: Serine to Cysteine at Amino Acid 680
S235F: Serine to Phenylalanine at Amino Acid 235
SARS-CoV-2: Severe Acute Respiratory Syndrome Coronavirus 2
SNP: Single Nucleotide Polymorphism
T1001I: Threonine to Isoleucine at Amino Acid 1001
T716I: Threonine to Isoleucine at Amino Acid 716
TBE: Tris/Borate/Ethylenediaminetetraacetic acid
Y73C: Tyrosine to Cysteine at Amino Acid 73
FDA: United State Food and Drug Administration
VOC: Variant of Concern
VTM: Viral Transport Media

**Table Supplementary: List of Abbreviations and Definitions Used in Text**

